# Expression of vimentin intermediate filaments in epithelial cells promotes cell migration and cell matrix interaction in 3D

**DOI:** 10.1101/2025.09.18.677142

**Authors:** Camille Rodriguez, Hyuntae Jeong, Jiwon Kim, Lily A. Cordner, Paul Cao, Suganya Sivagurunathan, Stephen A. Adam, Robert D. Goldman, Ian Y. Wong, Ming Guo

**Affiliations:** Department of Mechanical Engineering. Massachusetts Institute of Technology. 77 Massachusetts Avenue. Cambridge, MA 02139; School of Engineering. Brown University. 184 Hope St Box D. Providence RI 02912; Biomedical Engineering Graduate Program. Brown University. 184 Hope St Box D. Providence RI 02912; Computational Biology Core, Center for Computation and Visualization. Brown University. 180 George St, Providence RI 02906; Department of Cell and Developmental Biology, Northwestern University Feinberg School of Medicine, Chicago, IL, USA

## Abstract

During a variety of physiological and pathological processes, such as development, wound healing, and tumor progression, epithelial cells collectively invade into their surroundings. Vimentin intermediate filaments (VIFs) are often observed to play a role in the epithelial cells located at the margins of 2D cultures. However, their role in 3D collective cell behavior remains underexplored. Here, we investigate how induced vimentin expression affects 3D multicellular architecture and mechanics in luminal breast cancer cells (MCF-7) that ordinarily express keratin intermediate filaments only. We find that vimentin expression significantly alters 3D cell cluster morphology, inducing protrusions and increasing boundary fluctuations. Furthermore, cells in vimentin-expressing clusters show enhanced, more stochastic migration. In addition, these clusters exert stronger and localized traction forces on the surrounding matrix, indicating increased cell-matrix interactions. Transcriptomic analysis corroborates these biophysical findings, revealing upregulated gene expression for cell migration and matrix adhesion, and downregulated cell-cell adhesion genes. Our results demonstrate that VIFs are critical in modulating 3D multicellular collective morphology and dynamics, promoting invasive-like behavior by enhancing cell migration and cell-matrix interactions. These results provide fundamental insights into understanding tissue morphogenesis and disease progression.

## Introduction

Epithelial tissues disorganize and collectively invade the extracellular matrix (ECM) during embryonic development, wound healing, and tumor progression [1–5]. During this process, epithelial cells weaken their cell-cell adhesions and reorganize their cytoskeletal architecture, including F-actin, microtubules and cytoskeletal intermediate filaments [6]. To enable invasion, cells on the periphery generate actin-rich protrusions to mechanically or enzymatically remodel the surrounding ECM [7]. These peripheral cells are often found to undergo epithelial to mesenchymal transition (EMT), evidenced by expression of vimentin intermediate filaments (VIFs) [3, 8]. However, while F-actin and microtubules are commonly studied for their roles in cellular mechanobiology and migration [9–12], the functional role of VIFs in remains underexplored, particularly in the context of a cell collective in 3D [13–15].

Recent studies show that VIFs form a highly stretchable network in mesenchymal cells which together with other cytoskeletal elements provide protection against large deformations [16–21]. Moreover, it has been found in 2D cultures that VIFs modulate lamellipodia formation [22] and “leader cell” phenotypes [23], and act to orient traction stresses [24, 25]. Remarkably, microinjection of vimentin monomers or transfection with vimentin cDNA is sufficient for human breast epithelial MCF-7 cells (which ordinarily only expresses keratins) to elongate and migrate with a mesenchymal phenotype in 2D [26]. Subsequent work with a cumate-inducible promoter of vimentin expression in MCF-7 revealed increased intracellular stiffness and weakened cell-cell adhesion, as well as upregulation of EMT transcription factors such as TWIST1 [27]. In comparison, MCF-7 cells cultured in reconstituted basement membrane (e.g. Matrigel) ordinarily form multicellular clusters (“acini”) with limited invasiveness [28, 29] and undergo apoptosis in collagen I due to their inability to remodel the ECM (e.g. low expression of matrix metalloproteinase (MMP) [30]).

Here we study the functional role of VIFs in 3D collective cell behavior using human epithelial MCF-7 cells in which we can control vimentin gene expression and VIF network assembly in live cells. We find that expression of VIFs in MCF-7 cells significantly enhances cell migration in 3D cell clusters, likely by increasing cell-matrix interactions evidenced by increased cluster boundary fluctuations and traction forces exerted by the cell clusters. Furthermore, through transcriptomics we observe significant upregulation of the expression of genes associated with cell migration and matrix-adhesion, consistent with our biophysical results.

### Vimentin-induced MCF-7 cell clusters exhibit more irregular morphology

To study the effect of VIFs in 3D collective cell behavior, we use human breast epithelia cells, MCF-7s which are engineered to allow for controlled expression of vimentin. The MCF-7 cell line is cloned in the cumate-inducible vector and transfected to express vimentin protein [27]; within 3 to 5 days of exposure to cumate these cells form extensive VIF networks in live cells (Fig. S1). To study collective behavior in 3D we embed uninduced (no vimentin) and induced (wild type vimentin networks) MCF-7 cells in an interpenetrating network gel comprised of alginate (5 mg ml^-1^) and Matrigel (4 mg ml^-1^). This gel has a shear modulus of approximately 300 Pa to mimic the mechanical properties of a low malignant breast tissue microenvironment [3, 31, 32]. We seed single cells within this gel and allow them to grow and form multicellular clusters over 14 days. Around day 7, these clusters have an average of 10 to 12 cells with a maximum diameter less than 100 µm. Around day 14, there is a significant difference between uninduced (no vimentin) and induced (WT vimentin) clusters, with the induced clusters being larger. Uninduced clusters have an average of 32 cells per cluster with an average diameter of 120 µm, while induced cells have an average of 54 cells per cluster with an average diameter of 146 µm (Fig. S2).

To confirm the presence of vimentin post-induction, we immunostained for nuclei, F-actin, microtubules, and VIFs in 7 day old cell clusters (Fig. 1a). Notably, vimentin is not visible in the uninduced clusters, but VIFs occur throughout the entire induced cluster. The spatial organization of F-actin and microtubules is comparable between uninduced and induced clusters. Interestingly, there are striking morphological differences between induced and uninduced cell clusters. Uninduced MCF-7 cell clusters exhibit a spherical morphology with a relatively smooth periphery, consistent with previous literature [26, 33]. With induced vimentin, MCF-7 cell clusters exhibit more disorganized morphologies with sharp protrusions extending into the surrounding matrix (Fig. 1a, Fig. S3). To quantify the changes of cluster morphology over time, we first count the number of protrusions (as indicated by white arrow in Fig. S3; see details in Methods). We find that uninduced and induced clusters have similar number of protrusions on day 3; after day 7, induced clusters have a significantly higher protrusion count (Fig. 1b). This is consistent with previous 2D studies showing that VIFs modulate different protrusion formations such as lamellipodia [22] or invadopodia [34]. To further quantify these morphological differences, we compare the area moment of inertia of the clusters from brightfield images of their largest cross section against a fitted circle of the same boundary (see details in Methods). The resulting ratio is defined as C2D, with a value of 1 indicating the cluster’s boundary is a perfect circle and any value greater than 1 indicating deviation from its circular fit. We observe that uninduced and induced clusters both have similar C2D values on day 3. However, from day 7 onward induced clusters exhibit significantly larger C2D, confirming an increased deviation from a spherical morphology in cell clusters with WT vimentin (Fig. 1c). These results demonstrate that the presence of vimentin in the induced clusters results in a more irregular morphology as compared to uninduced clusters.

**Figure 1.**
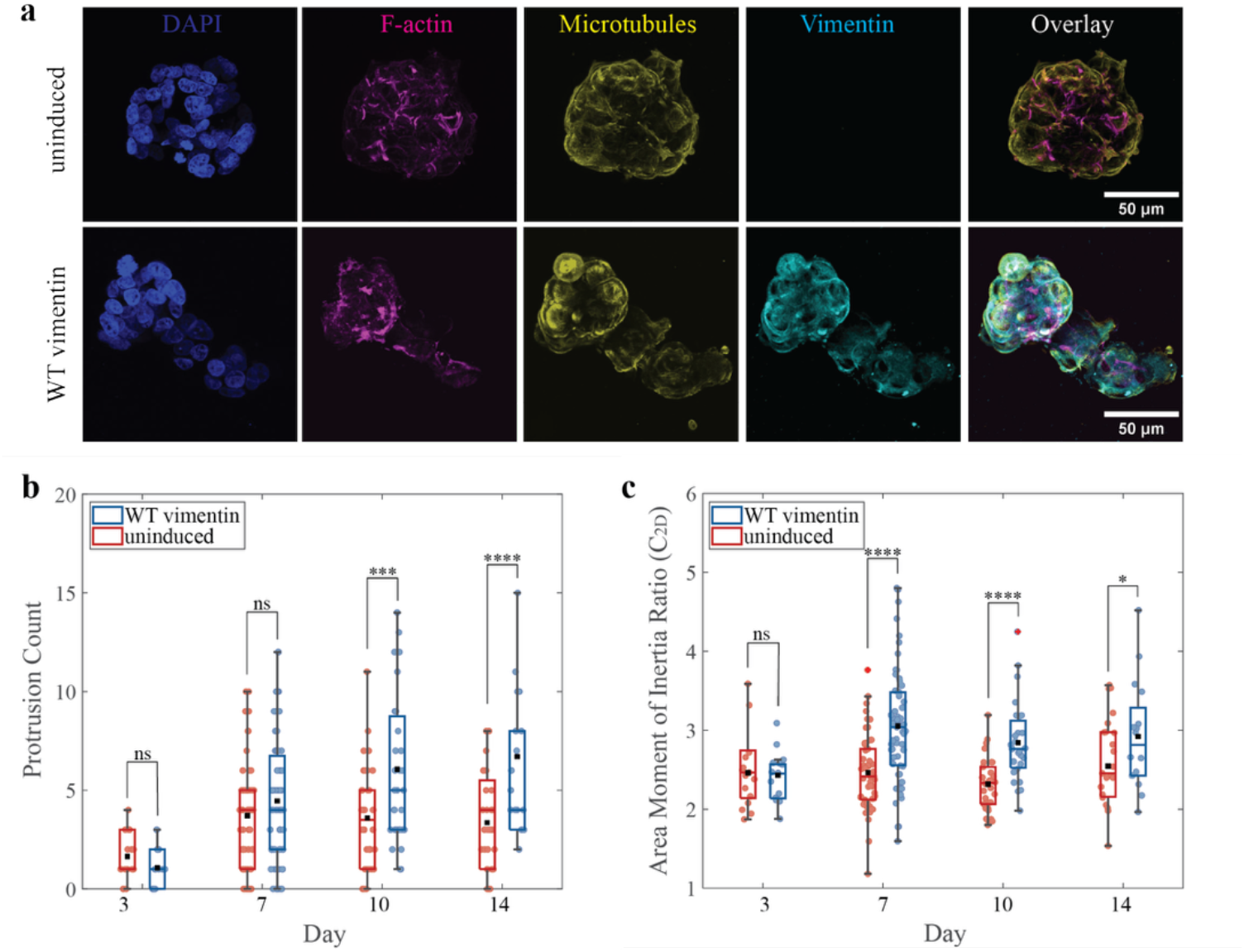
Morphological differences between uninduced (vimentin null) and induced (WT vimentin) MCF-7 clusters. **a**. Day 7 of cell clusters with the stained cellular components: nucleus, vimentin, microtubules, and F-actin. Uninduced clusters show minimal protrusions while induced clusters are more elongated with higher instances of protrusions. Details regarding definition of protrusions are in Fig. S3 and Methods. **b**. Protrusion count on each cluster at different days. On day 10 and beyond, there is a significant increase in protrusions of induced clusters (with WT vimentin) compared to the uninduced clusters (no vimentin). **c**. The area moment of inertia ratio (C_2D_) quantifies the morphology of the clusters. With increasing days, the induced clusters continue to deviate from their circular projection (with C_2D_ equals 1). On day 7 and beyond, the induced clusters have a significantly higher C_2D_ as compared to the uninduced clusters. Scale bar is 50 µm. The box plot in b and c represents the 25-75^th^ percentile of the data points, with the median (box line) and mean (black point) located within the box. The whiskers represent the minimum and maximum data points that fall within 1.5 times the interquartile range (IQR). N_b_ = 263, N_c_ = 253. * p < 0.1, *** p < 0.01, **** p < 0.001.

### Vimentin-induced MCF-7 clusters demonstrates dynamically fluctuating protrusions

The irregularity of cluster boundaries suggests potential changes in cell-matrix interactions upon expression of vimentin in MCF-7 cells. To investigate this behavior, we trace the temporal evolution of cluster boundaries over 12 hours at 10-minute intervals in order to evaluate the dynamic fluctuations of these boundaries. The uninduced cell clusters display weak fluctuations with a continual outward growth pattern. In contrast, the boundaries of the induced cell clusters (with WT vimentin) are much more dynamic; these clusters are constantly sending out and retracting their protrusions, as shown in Fig. 2a. The protrusions extend in multiple locations around the cluster. To measure the magnitude of the boundary fluctuations, we use the centroid of each cluster to obtain the relative change (δ) of the boundaries at various radial positions (θ) of the cluster between consecutive frames, as illustrated in Fig. 2b. To compare across clusters of various sizes, we normalize δ by the typical cell nuclear size of 10 µm found in our samples. We then compute the average magnitude of the normalized δ across all radial positions for consecutive frames; these results reveal that the induced clusters have significantly stronger boundary fluctuations (Fig. 2c). To further quantify the dynamics of these boundary fluctuations, we plot the normalized δ across time and angular position for the two representative clusters in Fig. 2a. In the uninduced cluster (no vimentin), we observe that boundary fluctuations are minimal as the normalized δ is near 0 throughout the entire 12 hours (Fig. 2d). With the presence of VIFs in the induced clusters, boundary fluctuations are significantly stronger with larger values of normalized δ values, as indicated by the blue and red colors in Fig. 2d. These fluctuations tend to occur at or near the same location and persist for several hours. These results reveal that with the presence of VIFs, cell clusters are more actively interacting with the matrix in 3D.

**Figure 2.**
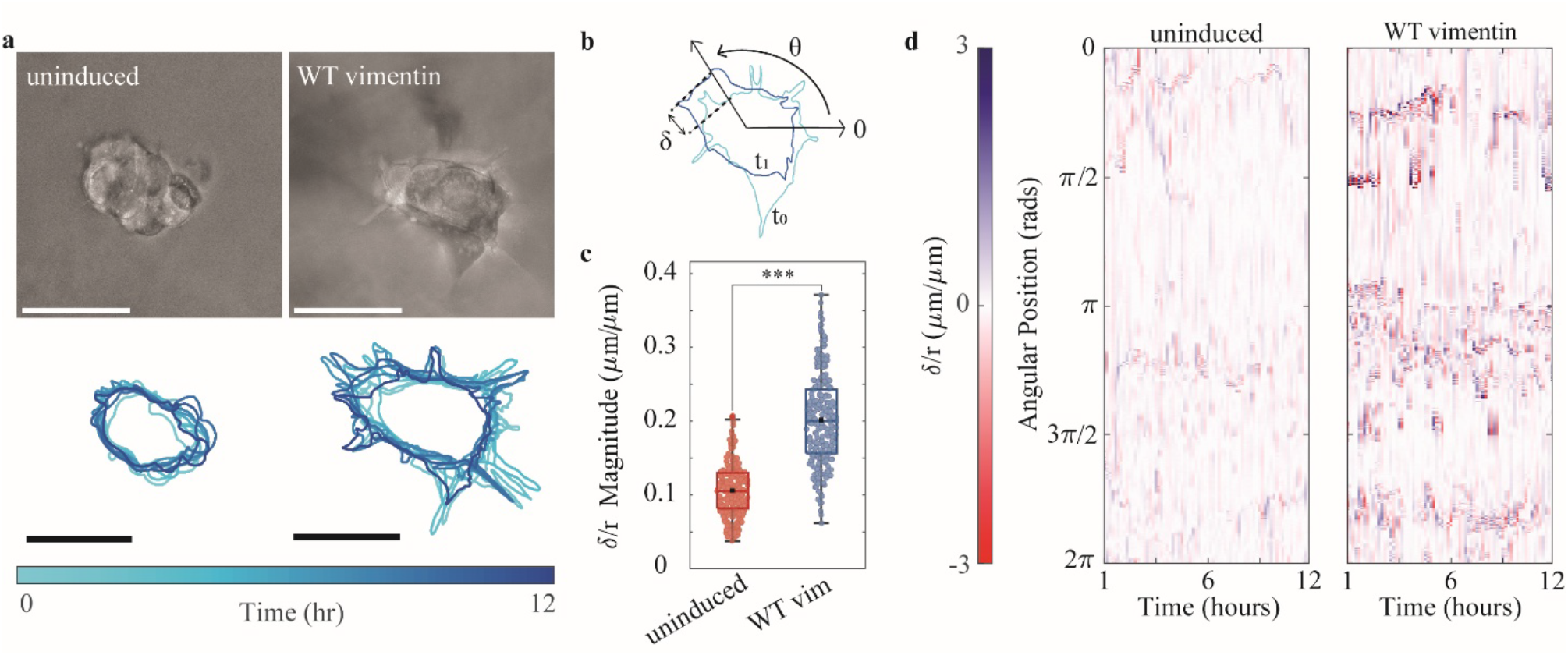
Spatial and temporal dynamics of boundary between cell cluster and hydrogel. **a**. From the brightfield images, the boundary between the cluster and the gel is outlined over 12 hours. The uninduced cluster’s entire body continues to grow outward whereas the induced cluster will extend and retract protrusions. **b**. The delta difference (δ) is the distance between two consecutive frames’ (t_0_ and t_1_) boundaries, along a radial position (θ) from the cluster’s centroid. **c**. Normalized over nuclear size (10 µm), the average magnitude of δ in between two frames is plotted for all samples. There is a significantly larger δ magnitude for the induced clusters. **d**. The δ magnitude of all 8 samples, averaged within each frame, exhibits large and repetitive protrusions with VIFs present. Scale bar is 50 µm. The box plot in c represents the 25-75th percentile of the data points, with the median (box line) and mean (black point) located within the box. The whiskers represent the minimum and maximum data points that fall within 1.5×IQR. N = 576. *** p < 0.01.

### Vimentin-induced MCF-7 clusters feature increased motility

Since the overall morphology of induced cell clusters (expressing WT vimentin) are more dynamic, we then investigate whether individual cells within the clusters are also more dynamic. To do so, we track individual cells in 3D and quantify their migratory trajectory and speed. The MCF-7 cells are transfected to express red fluorescent protein-tagged nuclear localization signal (RFP-NLS) which allows us to visualize individual cell nuclei. We image uninduced and induced clusters in 3D on day 7 and day 14 using confocal microscopy, over 12 hours at a 10-minute interval (Movies SI1 and SI2). To study cell migratory behavior, we track the position and movement of individual nuclei within the cluster using 2D projections. We find that there is a limited cell nuclear movement in the uninduced cell clusters as shown by short nuclear trajectories (Fig. 3a); in contrast, the cell nuclei of the induced cell clusters are significantly more dynamic and their trajectories traverse longer distances in comparable time, consistent with previous studies on 2D [26]. If we plot all the trajectories from their origin together, we observe that the trajectories of uninduced cells are indeed more localized whereas the trajectories of induced cells are more dispersed (Fig. 3a).

**Figure 3.**
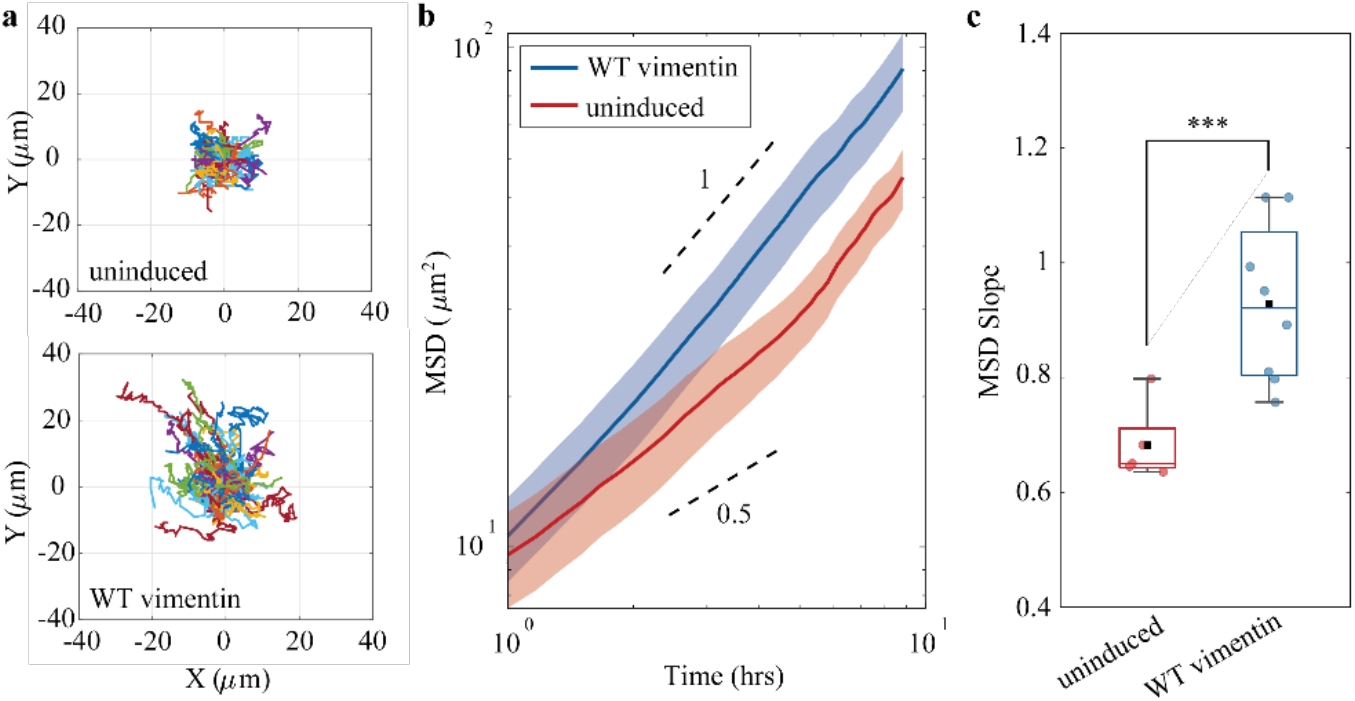
Migratory behavior of MCF-7 cells within 3D cell clusters. **a**. Trajectories of all cells’ nuclei (transfected to express RFP-NLS) within representative cell clusters (uninduced and induced respectively) are plotted together from their origin. Cells in uninduced clusters tend to stay near their starting position, while cells in induced clusters are able to move farther from their starting position within the cluster. **b**. The mean squared displacement (MSD) shows the induced cells have a significantly larger displacement over time. Represents the average MSD vs time of N_und_ = 5 and N_vim_ = 8 clusters. **c**. With an average slope nearing 1, the movement of nuclei of induced cells in clusters is consistent with random walk motion and increased speed as opposed to the more restricted migration of uninduced clusters. The box plot in c represents the 25-75th percentile of the data points, with the median (box line) and mean (black point) located within the box. The whiskers represent the minimum and maximum data points that fall within 1.5×IQR. N = 13. *** p < 0.01

To further quantify cell migration, we calculate the time and ensemble averaged mean square displacement (MSD) of all cells within each cluster, <Δ*r*^2^(τ)>, where Δ*r*(τ) = *r*(*t*+τ)-*r*(*t*). We observe that at long time scales (τ > 1 hr), cells in induced MCF-7 clusters have a significantly higher MSD, as shown in Fig. 3b. Moreover, we analyze the anomalous diffusion behavior by fitting the MSD vs time curve to a power law function (<Δ*r*^2^(τ)>=*βt*^*α*^). In this case, the exponent α here indicates the characteristics of the motion, i.e., diffusive (*α*=1), super-diffusive (*α*>1), or sub-diffusive (*α*<1). Interestingly, we find that uninduced MCF-7 cells show sub-diffusive behavior with *α*<1 (Fig. 3c). Sub-diffusive behavior typically arises from caging effects, and have been observed in dense granular systems [35, 36], as well as crowded 2D cell monolayers [37]. Here, both the high cell density due to the confined growth and cell-cell adhesion contribute to the intermittent caging effects. In contrast, we find that the cell migration in the induced MCF-7 cell clusters (with WT vimentin) become diffusive, as indicated by an average α ~ 1 (Fig. 3c). Previous studies have shown that vimentin expression leads to internalization of desmosomes and results in weaker cell-cell adhesion [26, 27]; this could lead to a weakened caging effect. Our results show that the presence of VIFs significantly changes cell migration characteristics in 3D cell clusters, potentially by regulating cell-cell adhesions.

### Vimentin-induced MCF-7 cell clusters further deform surrounding matrix

Multicellular clusters of MCF-7 cells with induced vimentin expression exhibited more dynamic protrusive morphologies as well as increased motility, suggesting mechanical differences in cell-matrix interaction relative to the uninduced clusters. To investigate cell matrix interaction, we employ 3D traction force microscopy to determine how cell-generated forces deform the surrounding 3D matrix [12, 38, 39]. To this end, we analyze the relative displacement of embedded fluorescent tracer particles (0.5 µm in diameter) in consecutive images at 10 min intervals (see details in Methods); matrix displacements toward the cell cluster indicate contractile forces generated by the cells within the cluster, while outward gel displacements indicate protrusive forces generated by cells pushing the gel. Consistent with the enhanced cluster boundary fluctuation and cell migration with vimentin expression, we find that induced MCF-7 cell clusters (WT vimentin) generate significantly greater matrix deformations, as compared to the uninduced ones (no vimentin); arrows mark amplitude and direction of local displacement of two representative cell clusters, uninduced (Fig. 4a, top) and induced (Fig. 4b, top) respectively. Induced MCF-7 clusters also exhibit sporadic “hotspots” of protrusion and contraction (~10 µm) that are localized to specific locations at the periphery (Fig. 1b, 20-30 min). To quantify this spatial non-uniformity, we bin the matrix displacements into equal regions from the centroid of the cluster and decompose them into the radial components [12, 40]. Indeed, both contractile and protrusive displacements are minimal for the uninduced cell clusters, as shown in Fig. 4a, bottom. With VIFs present in the induced cell clusters, we find that both contractile and protrusive displacements become significantly larger, reaching a mean displacement of ~2.5 µm over 10 min for corresponding bins (Fig. 4b, bottom). Interestingly, we observe that strong protrusions in one region may be immediately followed by strong contractions, consistent with the cyclical fluctuations observed in morphology (Fig. 2d). To further quantify the cell cluster generated displacements, we plot the probability density function (PDF) of displacement values around each cluster. Indeed, the PDF for uninduced clusters is much narrower (mainly near zero) compared to that for induced clusters (Fig. 4c). When compared across all samples, there is a significant increase in the number of higher displacement values, both contractile (0-5^th^ percentile) and protrusive (95-100^th^ percentile), in the induced clusters (Fig. 4d). Overall, these results demonstrate that the presence of VIFs promotes the mechanical interactions of cells and cell clusters with their surrounding matrix in 3D.

**Figure 4.**
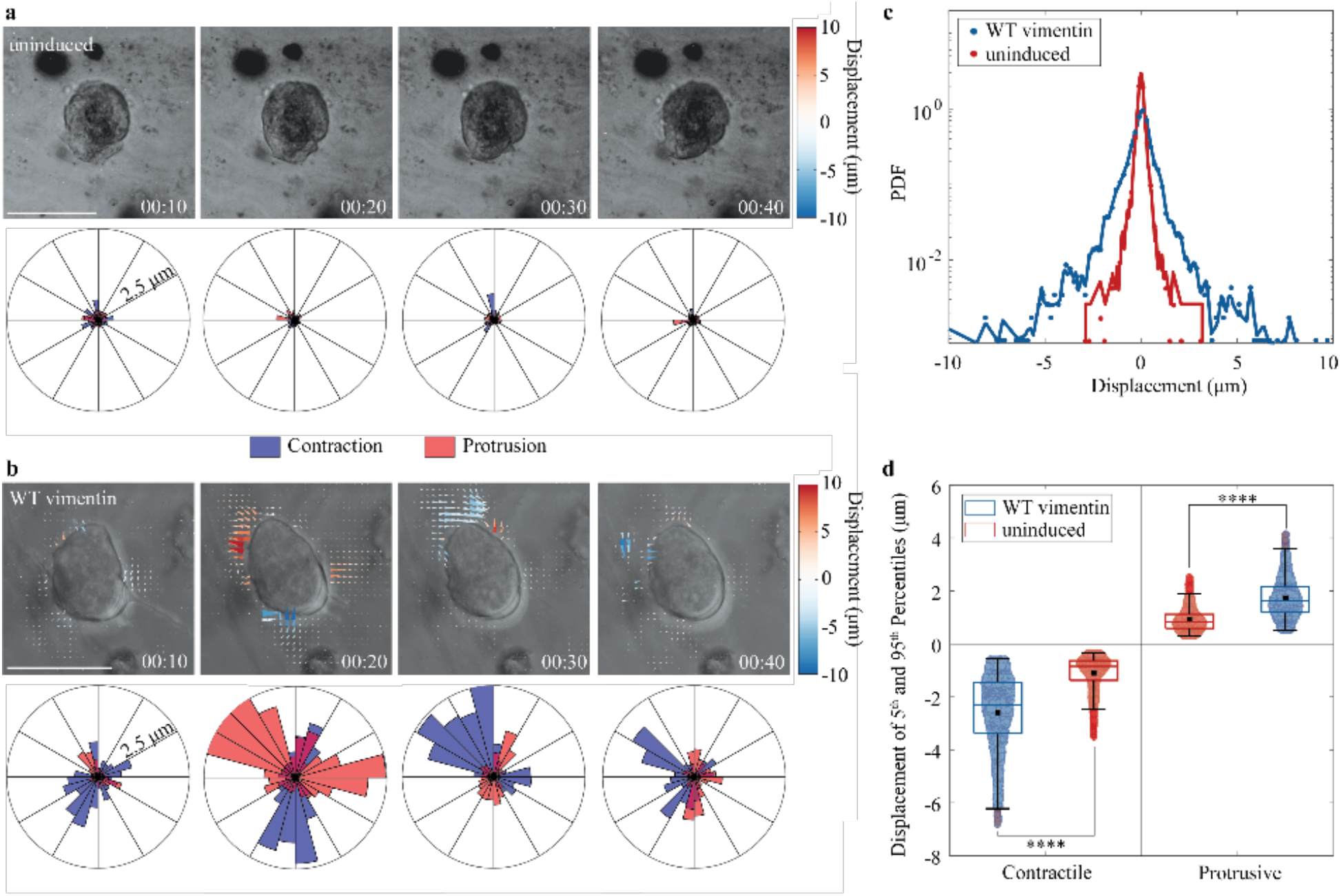
Hydrogel displacements generated by MCF-7 clusters. **a**. Displacement field between two consecutive time frames (interval of 10 minutes), displacement is determined by embedded particle movement. Uninduced clusters generate minimal gel displacements with the average in each sample being less than 2.5 µm. **b**. The induced clusters generate significantly larger gel displacements, both contractile (with arrows pointing towards the cluster) and protrusive (with arrows pointing away from the cluster). **c**. The probability density function examines the distribution of local gel displacements surrounding a cluster. The induced cluster exhibits a wider distribution indicating higher instances of large displacement magnitudes. Average PDF vs displacement of N_und_ = 5 and N_vim_ = 5 clusters. **d**. The highest and lowest 5% of all samples’ displacement fields shows that induced vimentin clusters have a wider distribution in both protrusive (positive displacement) and contractile (negative displacement) behavior. Scale bar is 50 µm. The box plot in c represents the 25-75th percentile of the data points, with the median (box line) and mean (black point) located within the box. The whiskers represent the minimum and maximum data points that fall within 1.5×IQR. N = 5259. **** p < 0.001

### Vimentin induction upregulates pro-migratory genes and suppresses cell-cell adhesion genes

To gain mechanistic insight into the regulation of the biophysical changes in the phenotype of clusters within 3D matrices, we analyze RNA-seq data previously acquired for both uninduced and induced MCF-7 cell lines grown in 2D cultures (GSE206724 [27]). We profile gene lists based on the 494 differentially expressed genes between vimentin induction relative to uninduced, using g:Profiler [41]. Notably, the top hits are terms associated with cell periphery, developmental process, cell migration, and cell adhesion (Fig. 5). In vimentin-induced clusters, cells upregulate genes associated with actin-based cell projections and a negative regulation of cell-cell adhesion genes including podocalyxin-like protein 1 (PODXL), a CD34-related sialomucin which weakens cell-cell adhesion [42]. Indeed, PODXL overexpression is sufficient to increase collective invasion of MCF-7 cells in clusters grown in 3D matrices [43], as well as the formation of protrusive invadopodia [44]. As a scaffolding protein PODXL facilitates interactions between the cell membrane and the cytoskeleton inducing migration by enhancing actin-rich lamellipodia protrusions, along with promoting budding of cohesive nodules [45]. Moreover, there is downregulation of the armadillo protein plakophilin 1 (PKP1) which links desmosomes to the keratin cytoskeleton [46]. Finally, there is upregulation of FAT1 cadherin and Roundabout1 (ROBO1), both are regulators of cell-cell adhesion and protrusive actin structures [47–49]. Thus, vimentin induction activates a coordinated program of gene expression that is consistent with our functional observations of increased cell protrusions and motility. Our analysis of gene expression further validates our experimental results as we observe induced clusters to be more dynamic inducing the expression of genes known to regulate increased migration, boundary fluctuations, and cell-matrix adhesion.

**Figure 5.**
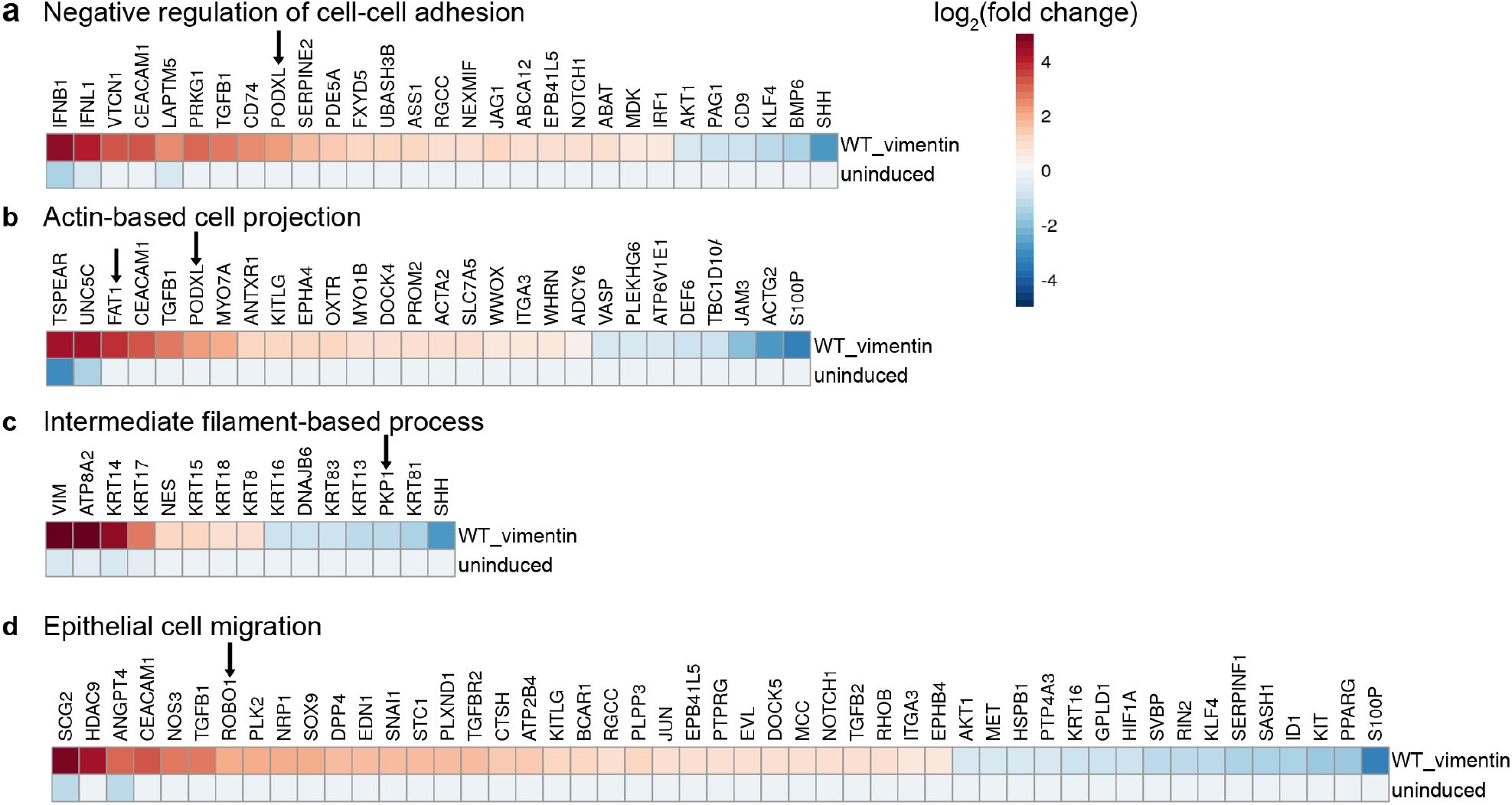
Comparison of gene expression of MCF-7 cells without or with induced vimentin expression. Heatmaps of gene expression associated with **(a)** negative regulation of cell-cel adhesion, **(b)** actin-based cell projection, **(c)** intermediate filament-based process and **(d)** epithelial cell migration.

## Conclusion

Despite the well-established finding that VIFs, a marker for the EMT and vimentin expression in the EMT has been linked to numerous physiological processes, and their well-documented role in regulating the mechanical properties of cells, their roles in the behavior of a 3D multicellular collectives remains largely unknown. By using human MCF-7 epithelial cells programmed to inducibly express vimentin in 3D cluster cultures, we find that the induction of vimentin expression significantly changes cluster morphology, cell migration, and cell matrix interaction. Consistent with previous 2D studies [26], we find that the induced cells with WT vimentin migrate faster and in a random fashion in 3D multicellular clusters, as compared to the uninduced cells without vimentin which migrate slower and in a confined way. In addition, we find that MCF-7 clusters with WT vimentin significantly deviate from a spherical morphology with dynamically fluctuating boundaries and develop evident protrusions penetrating into surrounding matrices. Indeed, using TFM, we observe that the induced MCF-7 clusters (with WT vimentin) generate larger displacements in the surrounding gel, both protrusive and contractile, as compared to the uninduced cell clusters, confirming increased cell-matrix interactions upon induction of vimentin. These observed biophysical alterations are supported by our transcriptomic analysis revealing the upregulation of cell migration related genes and the down regulation of cell-cell interaction related genes. Our results show that the presence of VIFs in cells significantly changes the morphology and dynamics of 3D multicellular collectives, enhancing cell migration and promoting invasive-like behavior through cell-matrix interactions. These findings contribute to a clearer understanding of how VIFs, as a key cytoskeletal element, influence collective cell behavior in 3D environments. Such insights are valuable for ongoing efforts to unravel the complexities of tissue morphogenesis during embryonic development, wound healing, and the invasive mechanisms of tumor progression.

## Acknowledgements

This work was funded by an NSF Graduate Research Fellowship (CR), NIGMS through R01GM140108 (CR, HJ, JK, RDG, IYW and MG) and P20GM109035 (LAC, PC, and IYW), as well as the Brown University School of Engineering through the Hibbitt Postdoctoral Fellowship (JK).

## Author Contributions

MG designed and supervised project. CR performed experiments; CR, HJ, JK, IYW and MG analyzed experimental data, LAC and PC analyzed gene expression; SS, SAA, and RDG provided the cell line; CR and MG wrote the manuscript with feedback from all authors.

## Notes

### Competing Interest Statement

The authors have declared no competing interest.

