## Supplemental Figures and Methods for "Expression of vimentin intermediate filaments in epithelial cells promotes cell migration and cell matrix interaction in 3D"

***Affiliation***


This file includes:

SI Figures

Captions of SI movies

Methods

### Supplementary Figures

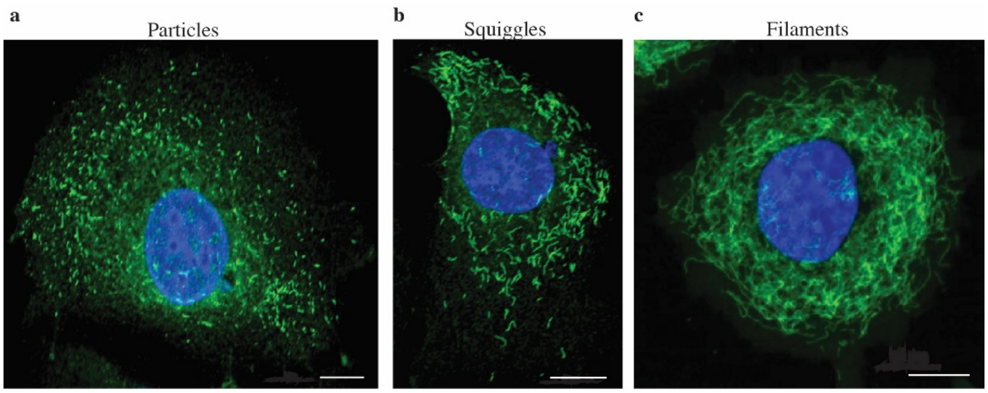


**Figure S1. Vimentin expression over multiple days in MCF-7 cumate-inducible cells a.** The expression of vimentin as particles (green) in MCF-7 on day 1 post cumate induction. Blue represents the nucleus of the MCF-7 cell. **b.** Vimentin progressed to squiggles on days 2 to 3 of post induction, indicating vimentin filament assembly. **c.** Vimentin forms an interpenetrating filamentous network on days 3 to 5 of post induction. Cells were fixed and stained with anti-vimentin. Scale bar is 10 μm.

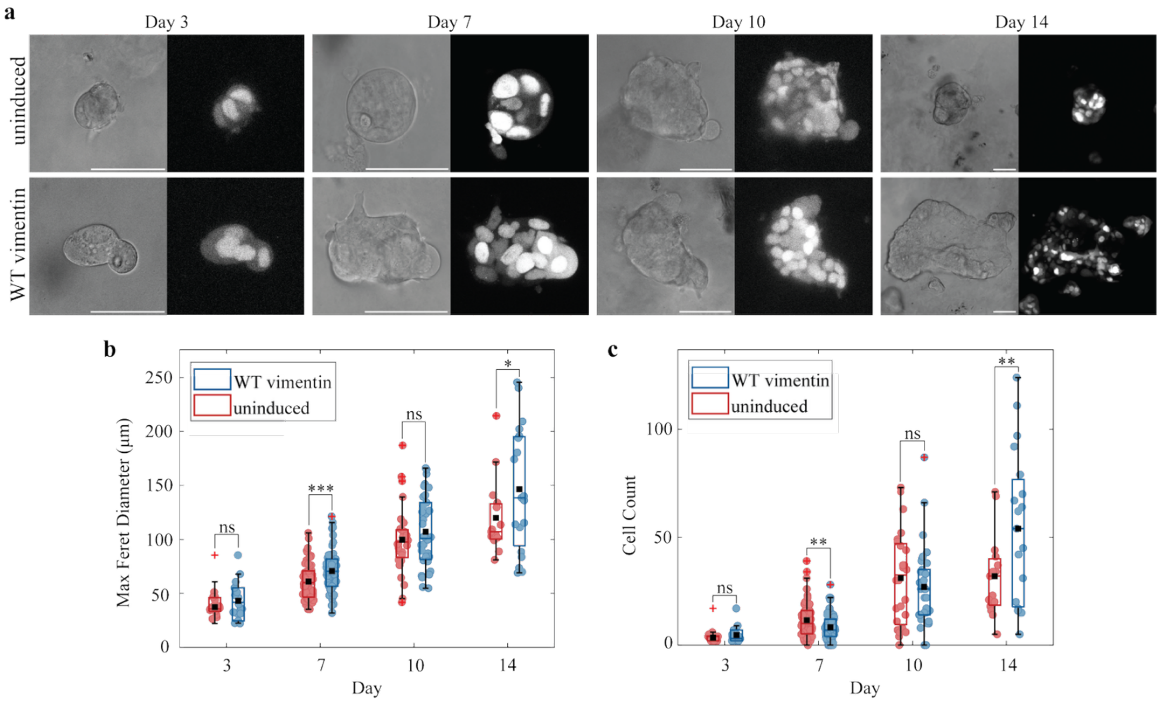


**Figure S2. Growth and size trends of uninduced and WT vimentin MCF-7 clusters over 14 days once embedded in the hydrogel. a.** Brightfield and fluorescent nuclei images of the growth progression of uninduced and WT vimentin (induced) MCF-7 cells at days 3, 7, 10 and 14. **b.** The maximum ferret diameter measured from the cluster's boundary outline, with an increase in both types of MCF-7s. **c.** The number of cells within each cluster is counted using fluorescent nuclei images, with the induced cell clusters exceeding the uninduced cell cluster on day 14. Scale bar is 50 µm. The box plot in b and c represents the 25-75$^{th}$ percentile of the data points, with the median (box line) and mean (black point) located within the box. The whiskers represent the minimum and maximum data points that fall within 1.5 times the interquartile range (IQR). N = 244. * $p < 0.1$, ** $p < 0.05$, *** $p < 0.01$.

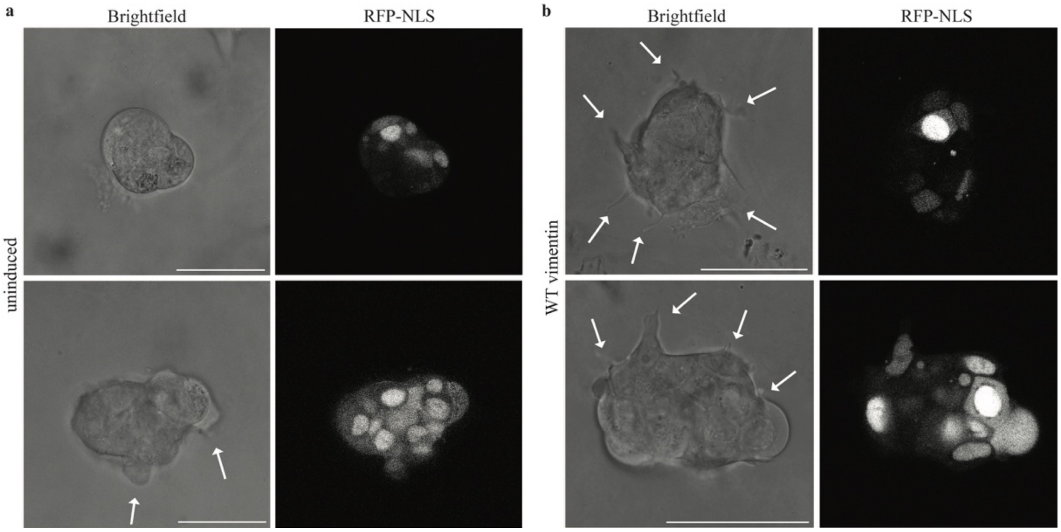


**Figure S3. Representative examples of both uninduced and WT vimentin clusters with types of protrusions examined. a.** Examples of uninduced clusters, one with no protrusions (top) and one with some of the smaller protrusions seen on these types of MCF-7s (bottom). **b.** Examples of WT vimentin (induced) clusters with multiple types of protrusions present, and one with one protrusion containing part of a nuclei (bottom). Scale bar is 50 µm. Arrows indicate protrusions.

**Movie S1. Uninduced cluster on day 7 showing the brightfield image and fluorescent nuclei of the MCF-7 cells. Scale bar is 50 µm.**

**Movie S2. WT vimentin cluster on day 7 showing the brightfield image and fluorescent nuclei of the MCF-7 cells. Scale bar is 50 µm.**

### Methods

#### 3D MCF7 cluster culture and hydrogel formation

**Cell Culture and Gel Preparation** To study the effect of VIFs in collective migration we use inducible MCF-7 cells that allow for controlled expression of vimentin. The MCF-7 cell line is cloned in the cumate-inducible vector and transfected to express vimentin proteins [1]. Within 3 to 5 days of exposure to cumate these cells form networks of vimentin (Fig. S1). To expand the study of collective behavior into 3D we use alginate-Matrigel hydrogel. Uninduced (no vimentin) and induced (wild type vimentin networks) MCF-7 cells are grown in the interpenetrating network gel comprised of 5 mg $ml^{-1}$ alginate and 4 mg $ml^{-1}$ Matrigel, with a shear modulus of approximately 300 Pa to mimic the mechanical properties of a low malignancy breast tissue microenvironment [2], [3]. Matrigel provides biological support for the cells while the alginate allows control over mechanical properties such as stiffness. The single cells in the gel then grow into multicellular structures form over 14 days. Around day 7 the clusters have approximately 5 to 40 cells with diameters ranging from around 50 to 110 μm. Around day 14 the clusters can have up to 125 cells and diameters up to around 250 μm (Fig. S2).

MCF-7 cells were stably transfected with a cumate-inducible vector for vimentin expression (pCDH-CuO-MCS-IRES-GFP-EF1α-CymR-T2A-Neo) [1] as well as mCherry-NLS for fluorescent labeling of the nucleus. Cells were kindly provided by Suganya Sivagurunathan and Stephen Adams (Northwestern University), and authenticated by STR profiling (ATCC). Cells are cultured in MEM supplemented with 10% fetal bovine serum (Invitrogen, 26140079), 1% MEM non-essential amino acids (Corning, 25-025-CI), 1% sodium pyruvate (Corning, 25-000-CI), 1% penicillin streptomycin, and 0.1% bovine insulin [1]. The cells are collected using 0.05% trypsin-EDTA solution (Corning, 45000-664) once they reach confluency in T-25 flasks in a normal cell culture incubator (37ºC with 5% $CO^2$). The cells grow without or without 5x cumate (inducer) (SystemBiosciences, QM150A-1) at least one week prior to being seeded in the gel. Of 100 µL of the collected cell suspension, 2 µL is added to the alginate-Matrigel mixture. The alginate-Matrigel interpenetrating network hydrogel is comprised of a final concentration of 5 mg $ml^{-1}$ alginate and 4 mg $ml^{-1}$ Matrigel (Corning,47743-706). To crosslink the gel $CaSO^4$ in PBS (no salt) (Corning,21-040-CV) is used with a final concentration of 0.043 mM in the gel. The gel precursor with seeded cells is placed in the incubator for 1 hour for gelation. After 1 hour, culture medium is added to the dish to keep the gel hydrated. Culture medium is then replaced every 3 days similar to normal MCF-7 culture procedures.

**Immunofluorescence staining** Immunofluorescence staining is done on the formed clusters on day 7 of growth in the gel. The 3D gels with the clusters are fixed with 4% paraformaldehyde for 30 minutes, permeabilized with 0.2% Triton X-100 (Sigma, T9284-500ML) diluted in PBS for 2 hours, and blocked with 0.5% bovine serum albumin (BSA) in PBS for 5 hours at room temperature. In between each step the gel was rinsed 3 times with PBS for 15 minutes each.

For vimentin intermediate filaments, cells are immunostained with vimentin polyclonal antibody (Invitrogen PA116759) in 1:400 dilution in PBS for 24 hours in 4ºC. Then Alexa Fluor 488 goat anti-chicken IgY (H+L) (Invitrogen A11039) in 1:400 dilution in PBS is added for 24 hours in 4ºC. The samples are protected against photobleaching from ambient light from this step on. For microtubules, the same steps were repeated except with alpha tubulin monoclonal antibody (Invitrogen A11126) then Alexa Fluor 568 goat anti-rat IgG (H+L) (Invitrogen A11077). For F-actin, Alexa Fluor 633 Phalloidin (Invitrogen A22284) in 1:200 dilution in PBS is applied and maintained for 24 hours 4ºC. For the nuclei, DAPI (ThermoFischer 62247) is diluted to a 300 nM solution in PBS was similarly maintained for 24 hours in 4ºC. In between each step the gel was rinsed 3 times with PBS for 15 minutes each. All images were recorded on a Leica DMI8 confocal microscope. Immunostaining images were taken in sequence to minimize wavelength crosstalk.

**Morphology measurements through microscopy**

MCF-7 clusters with RFP-NLS in the gels are maintained in a stage top live cell incubation chamber (5% $CO_2$, 37 °C) while imaging in 3D on a confocal microscope (Leica, TCL SP8). Both phase (cell boundaries) and fluorescence (RFP, cell nuclei) channels are imaged simultaneously. Static images of clusters are taken on days 3, 7, 10 and 14 after seeding in the hydrogel. Using a ×25/0.95 numerical aperture water objective, multiple z-stacks of a cluster are captured for morphology analysis. Separate samples were made for boundary fluctuations and cell migration analysis, to reduce cytotoxicity from multiple exposures or long time-scale imaging. On day 7 and 8, multiple z-stacks of a cluster were recorded every 10 minutes over 12 hours.

**Protrusion Count** Morphological analysis of clusters includes a protrusion count. Protrusions are counted at a magnification of 25x through the entire cluster's height. A minimum width of 0.5 µm with a minimum length of 1 µm are used as a preliminary dimension for a protrusion based on imaging quality and previous reported dimensions [4]. Additionally, it must have a length to width ratio less than 0.8 or greater than 1.3 to exclude the bulk body of the cluster. Examples of such protrusions can be seen in Fig. S3.

**Area Moment of Inertia** To capture additional deviations in morphology when comparing uninduced and induced clusters a ratio $C_{2D}$ was measured where $C_{2D} = \frac{I_b}{I_c}$ . The $C_{2D}$ uses the area moment of inertia of the cluster's outlined cell-matrix boundary ($I_b$) and compares it to a fitted circle on the same boundary ($I_c$). A $C_{2D}$ value of 1 would indicate that the cluster's boundary is a circle, as this value increases there is an increase in deviation from a circular fit. The phase channel was utilized in ImageJ to outline the cluster's boundary. The largest cross-sectional area of the cluster was used to calculate the area moment of inertia, using $I_b = \int r^2 dA$ . A fit ellipse is calculated in ImageJ from the cluster's boundary outline. Utilizing the major axis, we extrapolate an effective radius $R_{eff}$ to be used in the fitted circle area moment of $I_c = \frac{\pi R_{eff}^4}{4}$ .

**Cell-Matrix Interaction Analysis**

**Boundary Fluctuations** To quantify boundary fluctuations, we look at the temporal evolution of the cluster-matrix boundary over 12 hours at 10-minute intervals. From the phase channels the largest cross-sectional boundary is outlined and recorded in ImageJ. To quantify these changes, we measured the relative spatial change ($\delta$) between consecutive frames at regular radial positions (θ) around the centroid of the cluster. These values are then normalized by an average nuclear diameter of 10 µm to account for the various sizes of clusters across days.

**Cell Migration** Using a ×10/0.4 numerical aperture air objective, a 3D stack of clusters are captured to follow the cells' positions (through RFP-NLS) over 12 hours at 10-minute intervals. The fluorescent channel is projected into 2D for tracking the cells using TrackMate in ImageJ [5], [6]. The cell trajectories are zeroed by their starting position for plotting. From the trajectories we can calculate the displacement vectors and determine the mean squared displacement (MSD) between different time intervals.

**Traction Force Microscopy** To prepare the sample for traction force miscopy analysis, we modify some of the steps in cluster and gel formation. The cell suspension increased to 20 µL with an addition of carboxylate-modified polystyrene 0.5 µm fluorescent beads (Sigma, L3280-1ML). 3D image stacks are captured on day 7, 10, and 14 after seeding and gel formation using a ×25/0.95 numerical aperture water objective for 1 hour at 10-minute intervals. For analysis, about 3-4 slices of the midsection of the clusters are utilized due to clear x-y displacements seen in the boundary fluctuations. To ensure the fluorescent beads are captured in their entirety, each slice is a maximum 0.5 µm. These slices are then projected onto 2D for particle image velocimetry (PIV) based traction force microscopy using code previously established [7].

Displacement Arrays of Rendered Tractions (DART) analysis was performed on the 2D displacement data obtained from PIV analysis of tracer particles, inspired by our previous work [8], [9]. We constructed a displacement vector field $U_{grid}$, where $U_{grid}$ indicates the differences in the bead locations during 10 minutes. $U_{grid}$ was then decomposed into its radial, $U^{r}_{grid}$, and circumferential, $U^{\theta}_{grid}$, components based on the centroid points of spheroid masks.

$$U^{r}_{grid} = \left| U_{grid} \cdot n_{grid} \right| \tag{1}$$

$$U^{\theta}_{grid} = \left| U_{grid} - \left( U_{grid} \cdot n_{grid} \right) n_{grid} \right| \tag{2}$$

Subsequent classification of each displacement vector into protrusive and contractile was determined by the sign of $U_{grid}$. Each particle's position was binned into one of 24 angular regions depending on their location with respect to the center of the spheroid, and their displacement magnitudes classified according to their sign; (-: contraction, +: protrusion). The mean of each displacement type at each angular region was plotted to obtain a radial DART diagram. Here, the radius of DART diagram was set to 2.5 µm.

$$U^{i} = \begin{cases} U^{protrusive} & if\ U^{r}_{grid} > 0 \\ U^{contractile} & if\ U^{r}_{grid} < 0 \end{cases} \tag{3}$$

Utilizing a probability density function we then compare the displacements across time. All displacement points within the grid are fitted to a probability density function for one sample over 40 minutes, accounting for 5 consecutive frames. A kernel fit is utilized just for visualization purposes. From these distributions we examine the 5$^{th}$ and 95$^{th}$ percentile of each sample's distribution to examine the largest contractile and protrusive displacement values. Of these values, outliers are removed if these values are more than 1.5 of the interquartile range above the $75^{th}$ percentile and below the $25^{th}$ percentile. This allows us to compare multiple samples to have a representation of the largest displacement values found and prevalence of contractile or protrusive types of displacement.
